# *Gluconobacter oxydans* Knockout Collection Finds Improved Rare Earth Element Extraction

**DOI:** 10.1101/2021.07.11.451920

**Authors:** Alexa M. Schmitz, Brooke Pian, Sean Medin, Matthew C. Reid, Mingming Wu, Esteban Gazel, Buz Barstow

## Abstract

Rare earth elements (REE) are critical components of our technological society and essential for renewable energy technologies. Traditional thermochemical processes to extract REE from mineral ores or recycled materials are costly and environmentally harmful^1^, and thus more sustainable extraction methods require exploration. Bioleaching offers a promising alternative to conventional REE extraction^2–4^, and is already used to extract 5% of the world’s gold, and ≈ 15% of the world’s copper supply^5,6^. However, the performance of REE bioleaching lags far behind thermochemical processes^2,7–9^. Despite this, to the best of our knowledge no genetic engineering strategies have yet been used to enhance REE bioleaching, and little is known of the genetics that confer this capability. Here we build a whole genome knockout collection for *Gluconobacter oxydans* B58, one of the most promising organisms for REE bioleaching^10^, and use it to comprehensively characterize the genomics of REE bioleaching. In total, we find 304 genes that notably alter production of *G. oxydans’* acidic biolixiviant, including 165 that hold up under statistical comparison with wild-type. The two most impactful groups of genes involved in REE bioleaching have opposing influences on acid production and REE bioleaching. Disruption of genes underlying synthesis of the cofactor pyrroloquinoline quinone (PQQ) and the PQQ-dependent membrane-bound glucose dehydrogenase all but eliminates bioleaching. In contrast, disruption of the phosphate-specific transport system accelerates acid production and enhances bioleaching. We identified 6 disruption mutants, that increase bioleaching by at least 11%. Most significantly, disruption of *pstC*, encoding part of the phosphate-specific transporter, *pstSCAB*, enhances bioleaching by 18%. Taken together, these results give a comprehensive roadmap for engineering multiple sites in the genome of *G. oxydans* to further increase its bioleaching efficiency.

## Introduction

Rare earth elements (REE) are essential for the manufacturing of modern electronics^11–13^, and sustainable energy technologies including electric motors and wind turbine generators^14^; solid state lighting^15^; battery anodes^16^; high-temperature superconductors^17^; and high-strength lightweight alloys^18,19^. All of these applications place increasing demands on the global REE supply chain^20^. As the world demand for sustainable energy grows^21^, finding a reliable and sustainable source of REE is critical.

Current methods for refining REE often involve harsh chemicals, high temperatures, high pressures and generate a considerable amount of toxic waste^22^. These processes give sustainable energy technologies reliant on REE a high environmental and carbon footprint. As a consequence, due to its high environmental standards, the United States has no capacity to produce purified REE^23,24^.

There is growing interest in biological methods to supplement, if not completely replace traditional REE extraction and purification methods^2,8,25–27^. Biological extraction (bioleaching) is already used to extract 5% of the world’s gold^5,6^, and ≈ 15% of the world’s copper supply^5,6^ (in fact, Cu biomining in Chile alone accounts for 10% of the world’s Cu supply^28,29^).

The performance of REE-bioleaching lags behind thermochemical processes. For example, while thermochemical methods have 89-98% REE extraction efficiency from monazite ore^7,30^, *Aspergillus* species can only achieve ≈ 3-5%^2^. The acid-producing microbe *Gluconobacter oxydans* B58 can recover ≈ 50% of REE from FCC catalysts^10^. However, techno-economic analysis indicates that even this extraction efficiency is still not high enough for commercial application^8^.

Recent efforts to improve bioleaching have focused exclusively on process optimization^4^. To our knowledge, no genetic approaches have yet been taken for any bioleaching microbe^31^. With recent advances in tools for reading and writing genomes, genetic engineering is an attractive solution for enhancing bioleaching. However, applying these tools to non-model microorganisms like *G. oxydans* can be a significant challenge^19^. And while there have been some promising advances for editing *G. oxydans ‘* genome^32–37^, we do not yet know where to edit.

In the presence of glucose, *G. oxydans* secretes a biolixiviant rich in gluconic acid^10^. This is produced by periplasmic glucose oxidation by the pyrroloquinoline quinone (PQQ)-dependent membrane-bound glucose dehydrogenase (mGDH)^38^. The final pH of the biolixiviant is a major factor in REE bioleaching^10^. But, gluconic acid alone fails to explain bioleaching by *G. oxydans:* pure gluconic acid is far less effective at bioleaching than the biolixiviant produced by *G. oxydans*^10^. This means that even the most successful efforts to up-regulate mGDH activity and gluconic acid production are unlikely to take full advantage of *G. oxydans’* biolixiviant production capabilities.

To characterize the genome of *G. oxydans* and identify a comprehensive set of genes underlying its bioleaching capabilities, we built a carefully curated whole-genome knockout collection of single-gene transposon disruption mutants using Knockout Sudoku^39,40^ (**Fig. 1**). Final pH of the biolixiviant is a good predictor for bioleaching efficiency^10^, thus we used acidification as a proxy for bioleaching potential and have thoroughly screened the collection to identify mutants that differ in their ability to produce acidic biolixiviant (**Figs. 2** and **3**). Finally, we demonstrate that a single gene disruption - only one of several potential enhancement strategies - can already significantly improve *G. oxydans’* bioleaching capabilities (**Fig. 4**).

**Figure 1.**
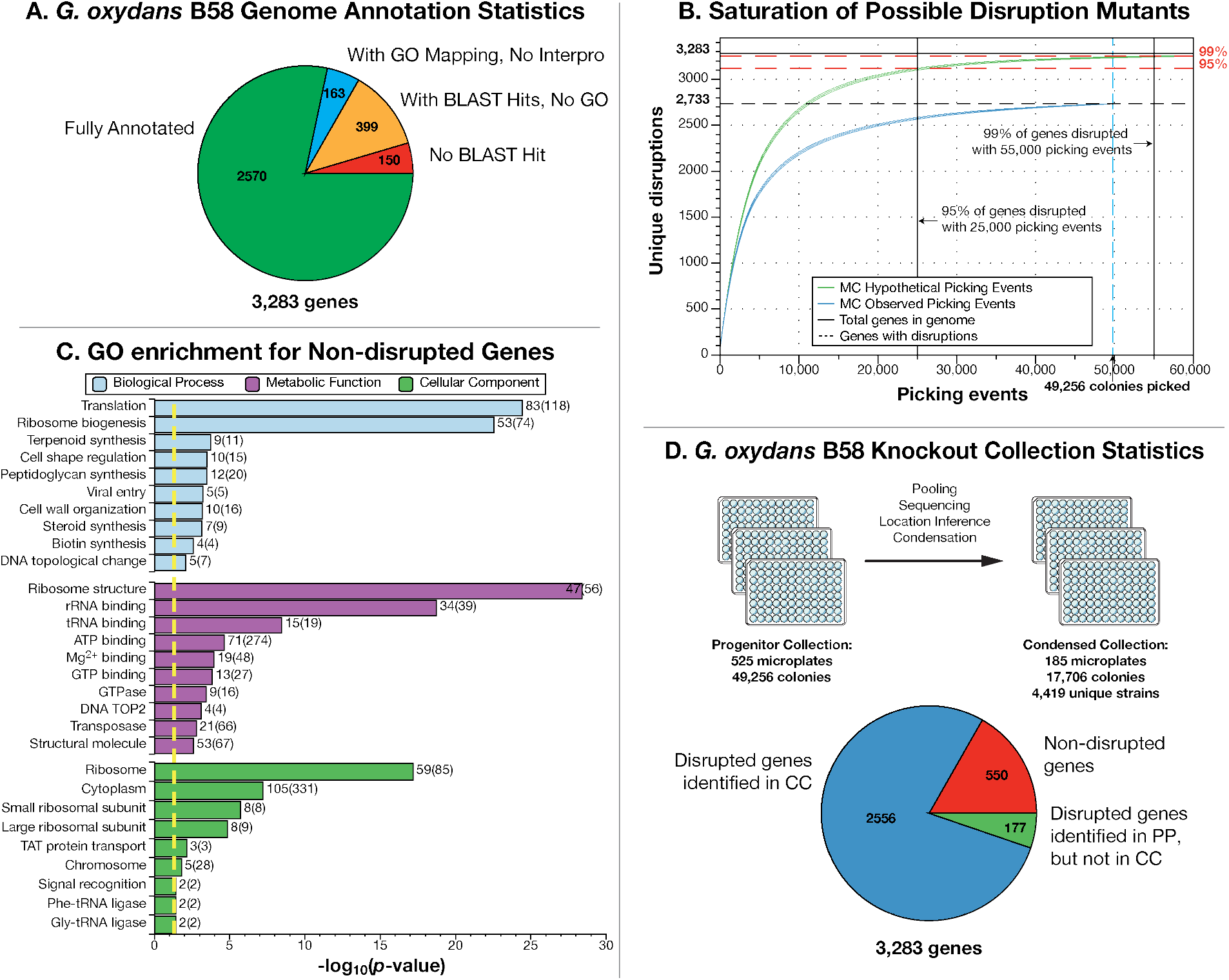
Knockout Sudoku was used to curate a saturating coverage transposon insertion mutant collection for *Gluconobacter oxydans* B58. (**A**) The *G. oxydans* B58 genome contains 3,283 genes. 2,570 genes were fully annotated with a BLAST hit, Interpro ID, and gene ontology (GO) group. An additional 163 genes have an annotation and GO group, but lack an Interpro ID, 399 only retrieved a BLAST hit, but no GO group, and 150 were unable to be assigned any annotation. (**B**) A Monte Carlo (MC) estimate of the number of genes represented by at least one mutant as a function of the number of mutants collected demonstrated that picking 25,000 mutants would yield at least one disruption for 95% of genes, while picking 50,000 mutants would yield at least one disruption for 99% of genes. In total, we picked 49,256 single-gene disruption mutants and located at least one disruption for 2,733 genes. A Monte Carlo simulation of picking with random drawing from the sequenced progenitor collection (PC) without replacements demonstrates that the genome coverage was truly saturated. The center of each curve is the mean value of the unique gene disruption count from 1,000 simulations while the upper and lower part of each curve represent two standard deviations around this mean. (**C**) A Fisher’s Exact Test for gene ontology enrichment among the non-disrupted (putatively essential) genes revealed significant enrichment (*p* < 0.05, yellow line) of genes involved in translation and other ribosome-related functions. (**D**) The curated condensed collection (CC) contains 17,706 isolated colonies across 185 plates. High-throughput sequencing of the CC confirmed the location for 4,419 unique disruption strains, representing disruptions in 2,556 genes. 177 genes located in the PC were not located in the CC. No disruption mutant was detected in 550 genes.

**Figure 2.**
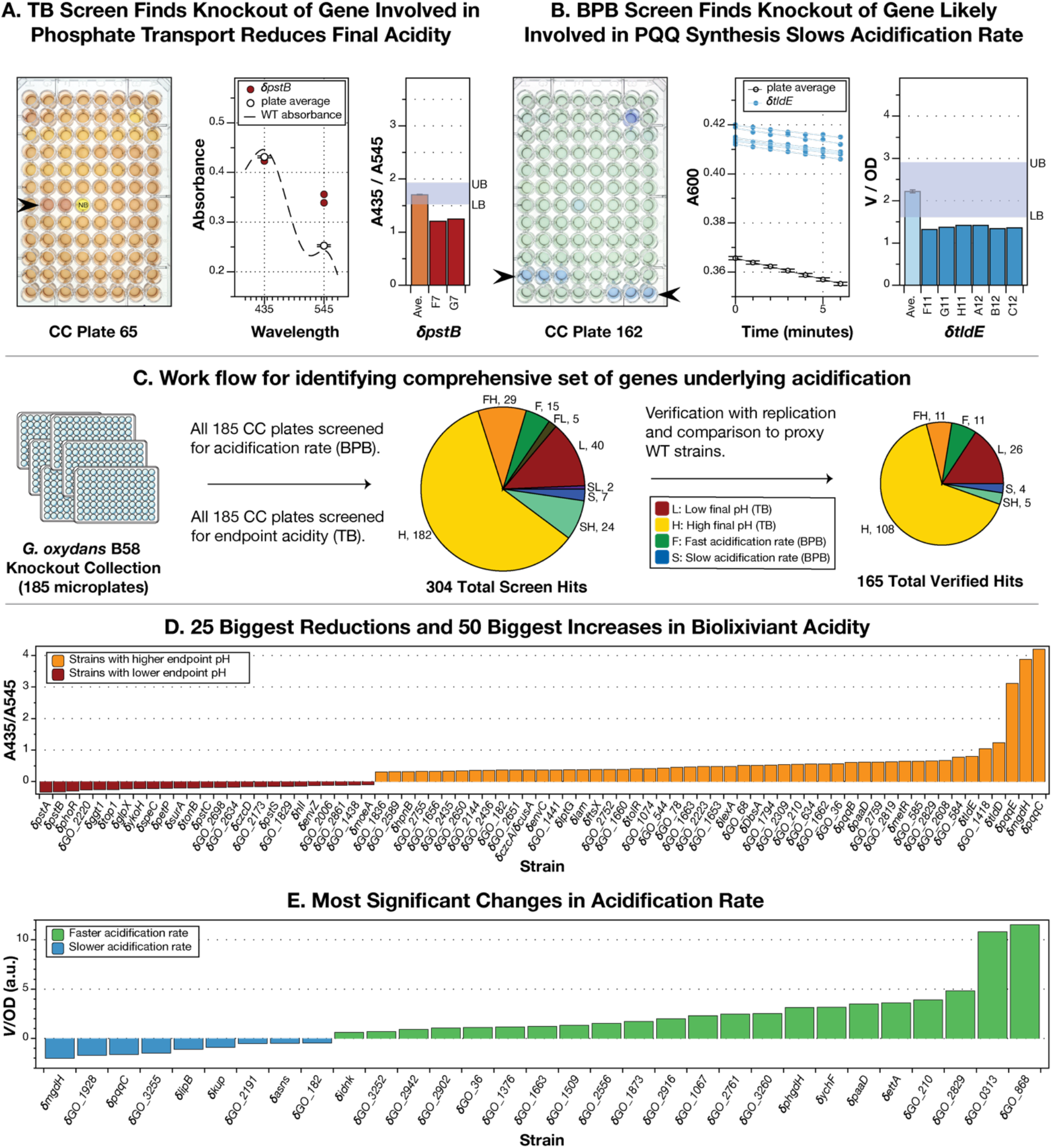
High-throughput pH screens of the *G. oxydans* whole genome knockout collection were used to identify genes that control REE bioleaching. (**A**) Thymol blue (TB) was used to measure the endpoint acidity of biolixiviant produced by each well of the condensed collection. The ratio of TB absorbance (A) at 435 and 545 nm is linearly related to pH between 2 and 3.4 (**Fig S1**). CC plate 65 contains biolixiviant produced by *δpstB* strain in wells F7 and G7 (arrowhead), whose absorbance at 435 nm and 545 nm is shown, along with the average absorbance of all wells on the plate. The dashed line represents a typical absorbance spectrum for WT-produced biolixiviant. The A435/A545 ratio for these two wells compared with the average ratio of the plate is well below the lower bound (LB) for the plate, indicating that *δpstB* produces a much more acidic biolixiviant than the average strain. (**B**) Bromophenol blue (BPB) was used to measure rate of change in pH at the onset of glucose conversion to organic acids. Rate was measured over a six minute period within five minutes of adding bacteria to a glucose and BPB solution. Condensed collection (CC) plate 162 contains the *δtldE* strain in wells F11 – C12 (arrowheads), whose changes in absorbance over time are graphed along with the average for that plate. A comparison of the normalized rate over OD for each well versus the plate average shows how *V*/OD for these wells was below the lower bound for CC plate 162. (**C**) All 185 plates of the CC were screened for acidification using the TB and BPB assays. Hits from both screens were verified in comparison with proxy WT strains. In total, 176 disruption strains were shown to significantly contribute to acidification by *t*-test with a Bonferroni-corrected alpha (α = 0.05 / # of comparisons). (**D**) The 25 largest reductions in biolixiviant pH, and 50 largest increases in biolixiviant pH (**Table S6E**). (**E**) All significant changes in acidification rate (**Table S6F**).

**Figure 3.**
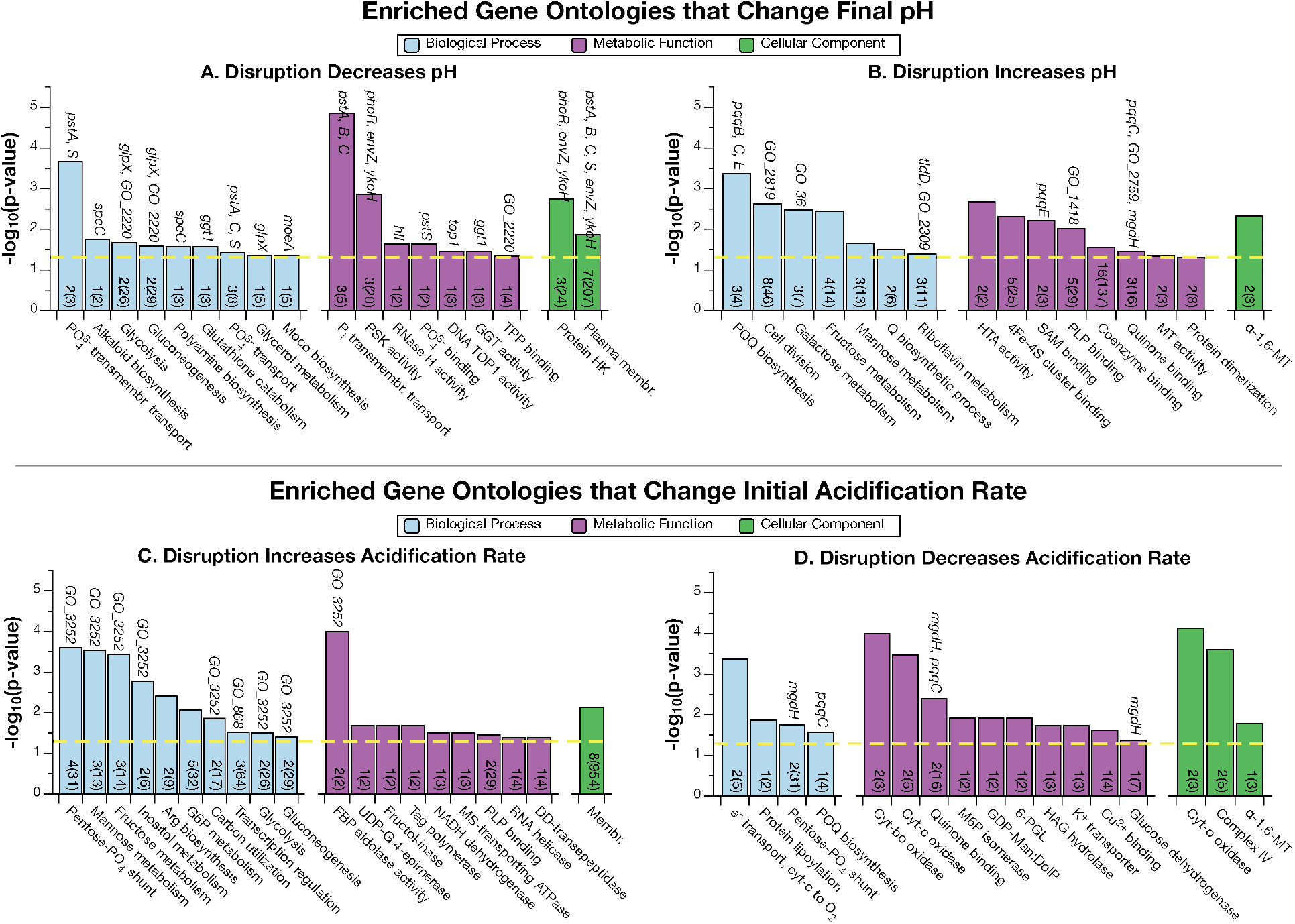
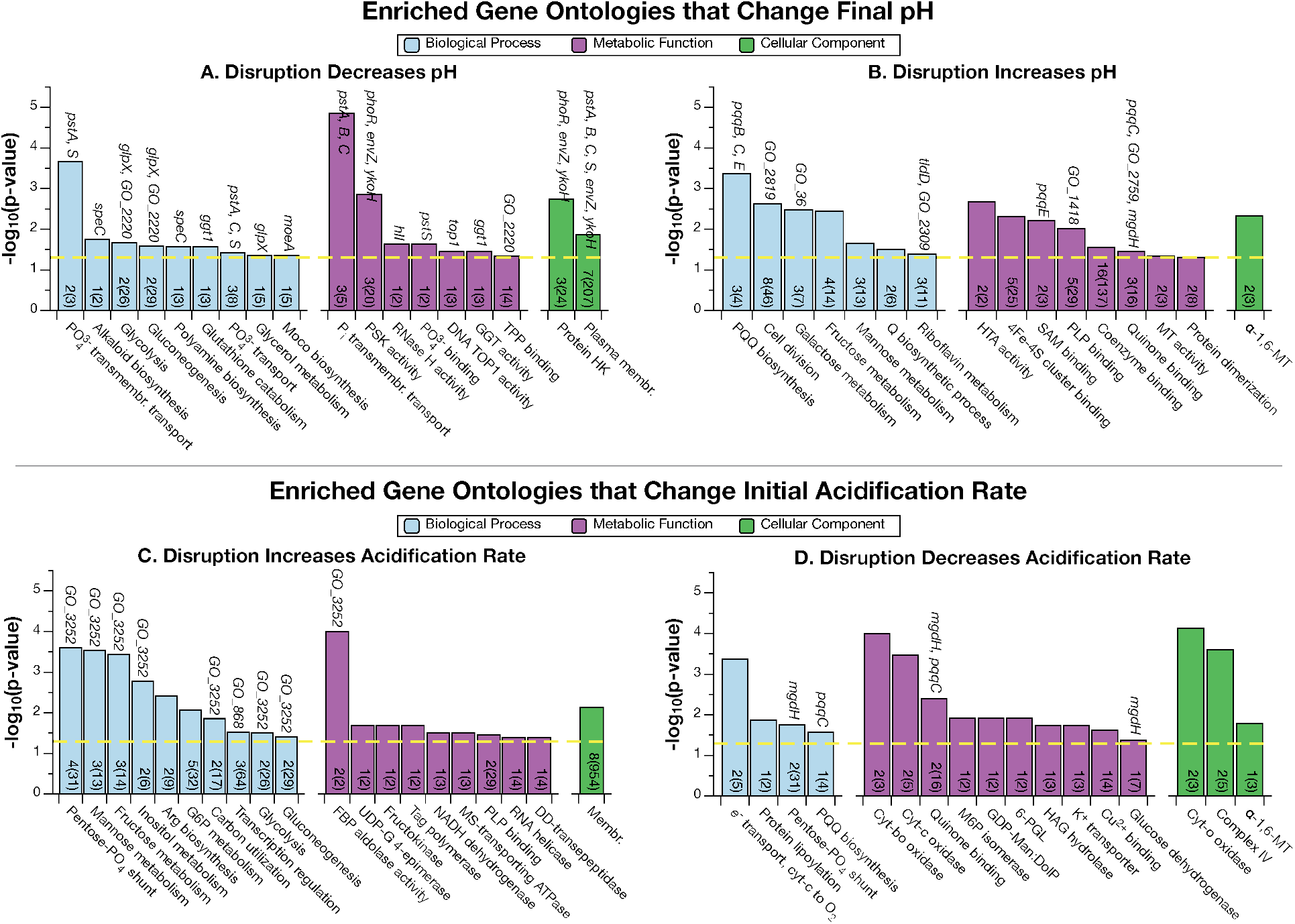
Genes involved in phosphate signaling, carbohydrate metabolism and PQQ synthesis were significantly overrepresented in the significant hits from high-throughput screens of acidification by *G. oxydans*. Fisher’s Exact Test was used to test for gene ontology enrichment (*p* < 0.05, yellow dashed line). Numbers at base of bars are how many genes from the significant hits are from that gene ontology (GO), out of the total in the genome (in parentheses). Genes selected for further analysis of endpoint pH and bioleaching (**Fig. 4**) that contribute to an enriched GO are listed above the bars. (**A** and **B**) Enriched GO among genes that decrease and increase end point pH. (**C** and **D**) Enriched GO among genes that increase and decrease initial acidification rate. Abbreviations: FBP: fructose-bisphosphate; GDP-Man:DolP: dolichyl-phosphate beta-D-mannosyltransferase; GGT: glutathione hydrolase; G6P: glucose 6-phosphate; HTA: homoserine O-acetyltransferase; DD-transepeptidase: D-Ala-D-Ala carboxypeptidase; HAG: hydroxyacylglutathione; Membr: membraneMoco: Mo-molybdopterin cofactor; MS: monosaccharide; MT: mannosyltransferase;M6P: mannose-6-phosphate; Pi: inorganic phosphate;PLP: pyridoxal phosphate;PQQ: pyrroloquinoline quinone;PSK: phosphorelay sensor kinase;Q: queuosine; RNase H: DNA-RNA hybrid ribonuclease;SAM: S-adenosyl-L-methionine; TPP: thiamine pyrophosphate; TOP1: topoisomerase type 1; HK: histidine kinase; UDP-G: uracil-diphosphate glucose; 6-PGL: 6-phosphogluconolactonase.

**Figure 4.**
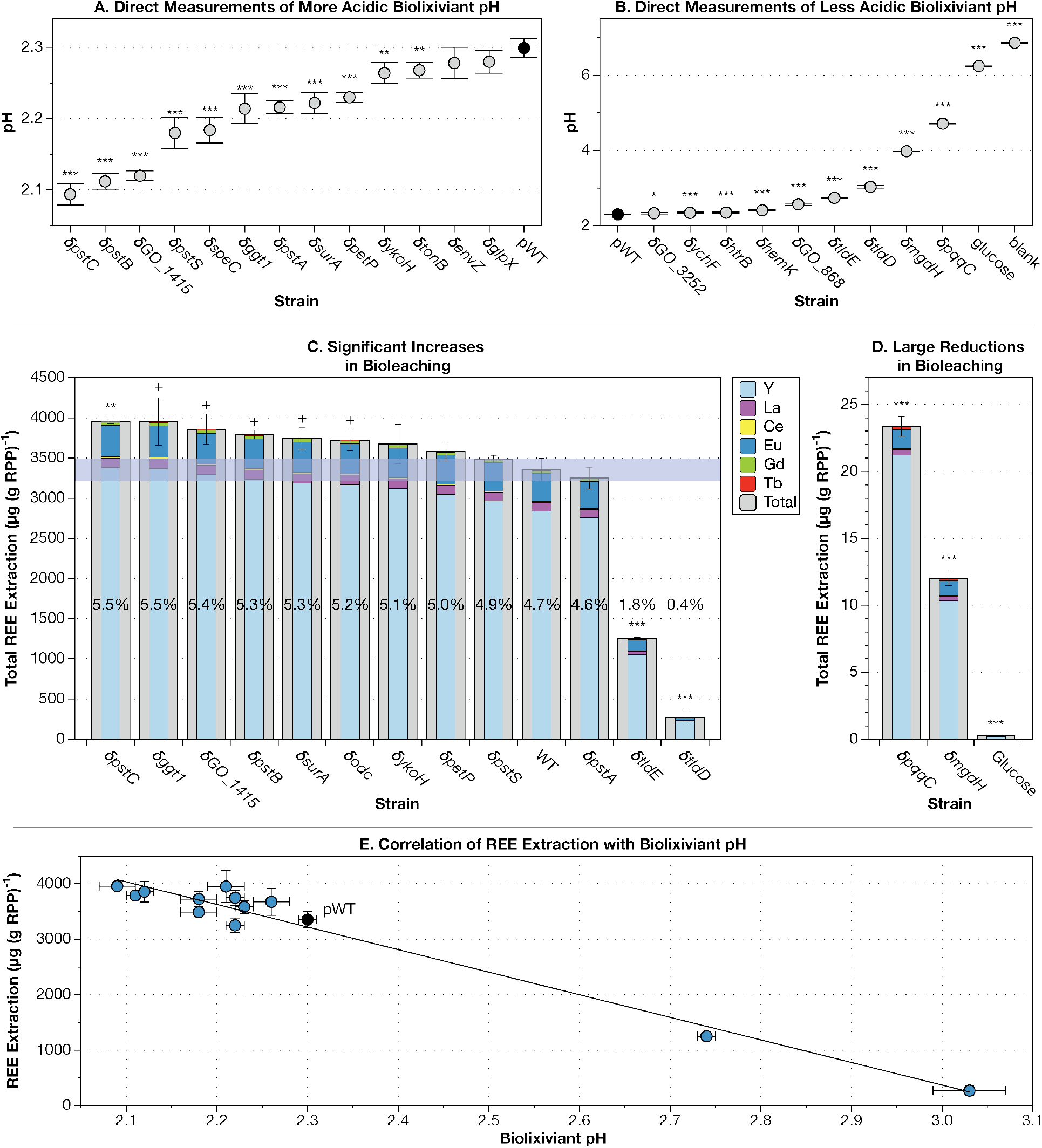
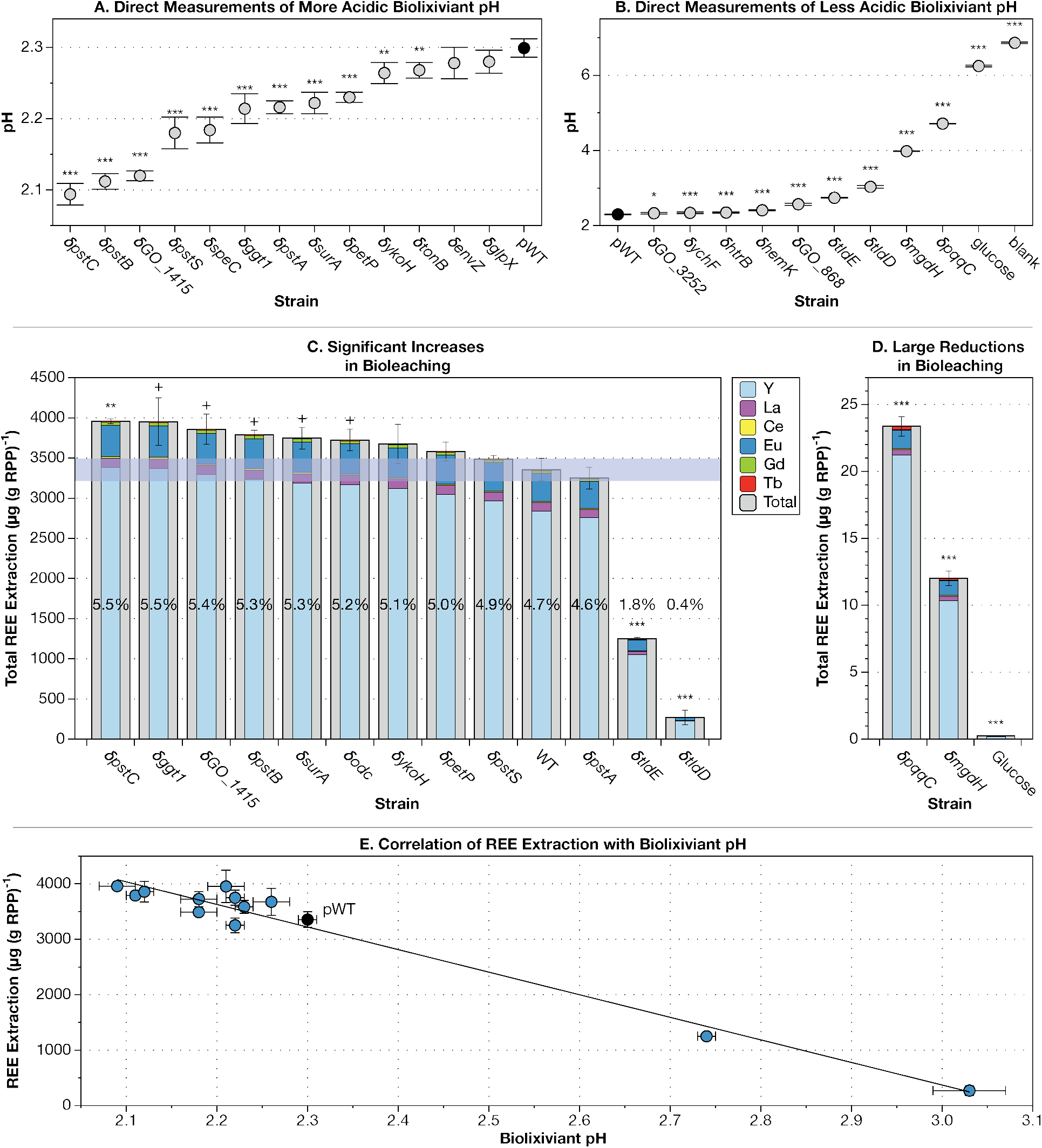
Increased acidification strains of *G. oxydans* B58 are able to increase rare earth extraction from retorted phosphor powder (RPP). (**A** and **B**) A subset of 20 disruption strains were tested for acidification with direct pH measurement. pH measurements significantly different from pWT (black circle) are labeled with asterisks: *, *p* < 0.05; **, *p* < 0.01; ***, *p* < 0.001 (n = 5, df = 18). Error bars represent standard deviation. (**C** and **D**) Ten disruption strains with the lowest final biolixiviant pH and four with the highest were tested for RPP bioleaching capabilities. Outer gray bars represent total REE extracted. Inner multicolored bars represent fractional contributions of each REE. Error bars represent standard error for total REE extracted. Percentages are total REE extraction efficiency (based on previously published REE amounts in the RPP [Reed2016a]). (**C**) Using a two-tailed *t*-test between each mutant and pWT demonstrated eight strains were significantly better or worse at bioleaching total REE (+,*p* < 0.05; n = 5, df = 18). With a Bonferonni correction, only one was significantly better (**, *p* < 0.01/12), but two of the higher pH biolixiviants that extracted detectable REE were significantly attenuated in bioleaching capability (* **,*p* < 0.001/12). (**D**) Disruption mutants for *mgdh* and *pqqC* are only able to extract less than 1% of the REE that wild-type *G. oxydans* can, but still extract significantly more REE than glucose alone when measured at a lesser dilution (***,*p* < 0.001/2). (**E**) Total REE extraction linearly correlates with pH. Error bars represent standard deviation for pH and standard error for total REE extracted.

## Results and Discussion

### Development of a Knockout Collection for *G. oxydans* Covering 2,733 Genes

We built a saturating coverage transposon insertion mutant collection for *G. oxydans* B58 and catalogued and condensed it with the Knockout Sudoku combinatorial pooling method^39,40^ (**Fig. 1**). We sequenced the *G. oxydans* B58 genome and identified 3,283 open reading frames (**Fig. 1A**; **Table S1**; **Materials and Methods**). Following the recommendation of Monte Carlo simulations, we collected 49,256 transposon insertion mutants (the progenitor collection; PC) to ensure saturating coverage of the *G. oxydans* B58 genome (**Fig. 1B**; **Materials and Methods**).

Progenitor collection sequencing results indicate that we were able to generate at least one disruption mutant for almost every non-essential gene in the *G. oxydans* genome. In total, we identified disruption strains for 2,733 genes out of the 3,283 genes in the *G. oxydans* B58 genome. Since every predicted gene contains at least seven AT or TA transposon insertion sites (**Table S1**), the remaining 550 non-disrupted genes are likely to be essential. A Fisher’s Exact Test for gene ontology (GO) enrichment representing 268 of the non-disrupted genes demonstrated significant enrichment (*p* < 0.05) in several essential ontologies, with the greatest enrichment in those relating to the ribosome and translation (**Fig. 1C** and **Table S2A**).

The progenitor collection catalog was used to create a condensed *G. oxydans* disruption collection with at least one representative per non-essential gene. 47 progenitor strains were verified by Sanger sequencing prior to condensing, of which 43 (92%) were confirmed to have the predicted transposon coordinate (**Table S3A**). We selected one mutant for all 2,733 disrupted genes, a second mutant for 2,354 genes, and a third mutant for 50 genes where mutant location information was poor. All mutants were struck out for single colonies, and 2-10 colonies per mutant were picked, depending on the predicted number of cross-contaminating disruption strains in the originating well. This condensed collection contains 17,706 mutants in 185 96-well plates (**Table S4**).

The condensed collection catalog was validated by a second round of combinatorial pooling and sequencing. Of the 17,706 wells in the condensed collection, we were able to confirm the identity for 15,257 (**Table S4**). We confirmed 25 of these wells by Sanger sequencing, and 100% have the predicted transposon coordinate (**Table S3B**). Among these wells, we were able to verify the identity and location of 4,419 independent transposon insertion sites, representing a disruption mutant for 2,556 unique genes (**Fig. 1D**). 1,587 genes are represented by more than one disruption, and 3,317 of all disruptions occur in the first half of the gene (**Table S4**).

### Genome-wide Screening Discovers 165 Genes Significantly Linked to Acid Production

We screened the new *G. oxydans* B58 whole genome knockout collection for disruption mutants with differential acidification capability (**Fig. 2**). We used the colorimetric pH sensitive dye Thymol Blue (TB) to screen for changes in final biolixiviant pH (**Figs. 2A** and **S1**), and Bromophenol Blue (BPB) to screen for changes in rate of acidification (**Fig. 2B**).

In total, we noted 304 genes that apparently controlled acidification (**Fig. 2C** and **Table S5**). The TB screen discovered 282 genes whose disruption leads to a differential change in biolixiviant acidity (**Fig. 2C** and **Table S5**). 47 mutants produced a more acidic biolixiviant, while 235 produced a less acidic one (**Fig. 2C** and **Table S5**). The BPB screen identified 82 gene disruptions with differential rate of acidification: 49 with a faster rate, and 33 with a slower rate. 60 mutants were identified by both screens (**Fig. 2C** and **Table S5**).

Overall, we identified 165 genes that significantly (*p* < 0.05) changed the final biolixiviant pH, rate of acidification, or both, but did not change the growth rate (**Fig. 2C** and **Table S6**). We re-arrayed disruption strains with differential acidification into new 96-well plates alongside proxy wild-type (pWT) strains (**Materials and Methods**) that have a transposon insertion in an intergenic region and show non-differential growth or biolixiviant production (**Fig. S2**). The new collection was re-assayed with the TB and BPB assays and the strength and significance of each result was determined by comparison with pWT through a Bonferroni-corrected *t*-test (**Materials and Methods**). Mutants that cause the 25 largest reductions, and 50 largest increases in endpoint acidity yet do not affect growth rate are shown in **Fig. 2D**. A full set of mutants that cause significant changes in acidification are listed in **Table S6**.

31 mutants that cause significant changes in acidification rate without changing growth rate are shown in **Fig. 2E** and **Table S6F**. However, 14 of the faster strains, including *δGO_868*, a disruption of a LacI type transcriptional repressor which was the fastest strain, produced a less acidic biolixiviant than the wild-type, indicating that targeting these genes for engineering a faster acidifier would likely be at the expense of a more acidic biolixiviant. None of the strains with a faster rate of acidification also created a more acidic biolixiviant. This result suggests that multiple genetic engineering interventions will be needed to construct a strain of *G. oxydans* that simultaneously produces a more acidic biolixiviant than the wild-type at a faster initial rate.

### Phosphate Transport and PQQ Synthesis are the Biggest Controllers of Acidification

We used a Fisher’s Exact Test to determine which biological processes, metabolic functions, and cellular components are enriched among the gene disruptions that significantly change biolixiviant acidification (**Fig. 3**). Among the disrupted genes that led to a stronger acidity, the most significant enrichment for all three GO categories involves the phosphate-specific transport system, represented by *pstA*, *pstB*, *pstC*, *pstS*, and*phoR* (**Fig. 3A**). Other enriched ontologies include those related to phosphate signaling and binding.

Among the disrupted genes that led to a weaker acidity, several enriched GO are related to the synthesis or use of the redox cofactor, PQQ, represented by*pqqB,pqqC,pqqE, tldD*, and *mgdh* (**Fig. 3B**). Other enriched ontologies include those related to carbohydrate metabolism.

Acidification rate is controlled by carbohydrate metabolism and respiration. Disruptions in the pentose phosphate pathway increase acidification rate (**Fig. 3C**). Meanwhile, disruptions of the electron transport pathway components are the most significantly enriched group of mutants that decrease acidification rate (**Fig. 3D**).

### Single Gene Knockout Mutants Can Significantly Change REE Bioleaching

#### Validation of Dye Assays by Direct pH Measurements

We selected 22 strains from those with greatest increase or decrease in acidification (**Figs. 2D** and **E**) for further testing. Dye pH measurements were validated by direct pH measurements. 11 of the 13 strains that produced significantly lower acidity biolixiviant in TB assays did the same in direct pH measurements (**Fig. 4A**). The most acidic biolixiviant was produced by a disruption in the phosphate transport gene, *δpstC* at pH 2.09. In fact, 4 of the 11 mutant strains that produced a more acidic biolixiviant were disrupted in genes involved in the phosphate-specific transport system (*δpstA, δpstB, δpstC*, and *δpstS*).

Additional disruptions that led to a more acidic biolixiviant included those in a hypothetical protein with no similarity to anything previously characterized (*δGO_1415*); a gamma-glutamyltranspeptidase (*δggt1*); a periplasmic chaperone (*δsurA*); an HTH-type transcriptional regulator (*δpetP*); a two-component system sensor histidine kinase (*δykoH*); a Pyridoxal 5-phosphate (PLP)-dependent ornithine decarboxylase (*δspeC*); and a TPR domain protein that is a putative component of the TonB iron uptake system (*δtonB*).

9 of the tested strains produced biolixiviant significantly higher in pH than pWT (**Fig. 4B**). The most alkaline biolixiviant was produced by a disruption in the PQQ synthesis system, *δpqqC*, at pH 4.71. While *δpqqC* produced very little acid, this is below the pH of glucose in media alone, indicating that some bacterial acidification still occurred. In fact, 3 of the 9 mutants that produce reduced acidity biolixiviant either synthesize PQQ (*δpqqC* and *δtldD*), or use it as a cofactor (*δmgdh*).

Additional disruptions that led to a more alkaline biolixiviant than pWT include a Fructose-bisphosphate aldolase class II (*δGO_3252*); a GTP and nucleic acid binding protein (*δychF*); a lipid A biosynthesis protein (*δhtrB*); a peptide chain release factor (*δhemK*); the LacI type transcriptional repressor that increases initial acidification rate (*δGO_868*); and the second component of a proteolytic complex with TldD (*δtldE*).

#### Disrupting the Phosphate Transport System Significantly Increases Bioleaching

We tested if 10 of the mutants that produced a more acidic biolixiviant could bioleach REE from retorted phosphor powder (RPP) from spent fluorescent lightbulbs more efficiently than pWT (**Fig. 4C**). For each mutant, the elemental composition of REE leachate was similar to that previously reported^10^. Six of these mutants significantly increased bioleaching. Two of the better bioleaching mutants disrupted the *pst* phosphate transport system (*δpstC* and *δpstB*). Overall, we found that bioleaching efficiency correlates with biolixiviant pH, as expected (**Fig. 4E**).

The *δpstC* mutant produced the most acidic biolixivant and extracted the most REE from RPP: 5.5% total extraction efficiency as compared with pWT’s 4.7%. Put another way, *δpstC* removed 18% more REE from RPP than pWT. This increase in REE extraction remains significant even after adjusting the α to account for the possibility of a difference due to chance alone (Bonferonni correction, see **Materials and Methods**). Without the adjustment, six of the better acidifiers were also better bioleachers than pWT (**Fig. 4C**). The remaining better bioleachers increased REE extraction by between 11% (*δodc*) and 18% (*δggt1*) (**Fig. 4C**). We speculate that disrupting phosphate transport and signaling de-represses acid production in *G. oxydans*. Six of the disruption strains that resulted in a lower biolixiviant pH (*δpstC, δpstB, δggt1, δpstA, δpstS*, and *δykoH)*, including three that increased bioleaching (*δpstC, δpstB, δggt1)*, along with many more identified by acidification high-throughput assays (*e.g., δphoR, δenvZ*), are involved in phosphate transport, sensing and signaling.

In its natural environment, *G. oxydans* produces biolixiviants to liberate phosphate from minerals, not metals^41–43^. Under phosphate-limiting conditions, the PstSCAB phosphate transporter will activate the histidine kinase, PhoR, which in turn phosphorylates the transcription factor PhoB, and activates the *pho* regulon, enabling phosphate solubilization and uptake^44^. Under sufficient phosphate conditions, PhoB is deactivated by PhoR, which in turn inhibits expression of genes involved in the phosphate-starvation response. We speculate that by disrupting *pstSCAB* or *PhoR*, we prevent *G. oxydans* from sensing when there is adequate phosphate in its environment and when to stop producing biolixiviants.

#### Disrupting *mgdh* and PQQ Synthesis Genes Significantly Decreases Bioleaching

We also tested REE extraction by 4 mutants that produce a less acidic biolixiviant than pWT, and all were worse bioleachers than pWT (**Figs. 4C** and **D**). Unsurprisingly, *δmgdh* mutant was the worst bioleacher of all tested, considering its lack of gluconic-acid production^38^. *δmgdh* reduced bioleaching by 97%. Disruption mutants that knocked out synthesis of mGDH’s essential redox cofactor, PQQ, also produced significant reductions in biolixiviant acidity. *δpqqC* reduced bioleaching by ≈ 94%. While bioleaching by *δmgdh* and *δpqqC* was negligible compared to pWT, they were able to bioleach a statistically significant amount of REE compared to glucose alone. This indicates, as previously speculated^10^, that a bioleaching mechanism independent of mGDH exists in *G. oxydans* (**Fig. 4D**).

Disruption mutants in *tldD* and *tldE* were also much worse at bioleaching than pWT. *δtldD* reduces bioleaching by 92%, while *δtldE* reduces it by 63% (**Fig. 4C**). We speculate that TldD and TldE contribute to the supply of the PQQ cofactor to mGDH. *δtldD* strongly attenuates acid production (**Fig. 4B**), and the gene has already been implicated in PQQ synthesis in *G. oxydans* 621H^45^. In *E. coli*, TldD and TldE form a two-component protease for the final cleavage step in the processing of the peptide antibiotic, Microcin B17^46^. In a similar manner, PqqF and PqqG from *Methylorubrum extorquens* form a protease that rapidly cleaves PqqA, the peptide precursor to PQQ^47^. We speculate that TldD in *G. oxydans* plays the same role as PqqF from *M. extorquens*, while TldE plays the same role as PqqG. Deletion of*pqqF* in *M. extorquens* completely inhibits cleavage of PqqA, while we find that disruption of *tldD* in *G. oxydans* reduces REE bioleaching by 92%. Moreover, deletion of*pqqG* in*M. extorquens* only reduces PqqA cleavage by 50%^47^, while disruption of *tldE* only reduces REE bioleaching by 63%. These parallels strongly indicate a novel role for TldE in the biosynthesis of PQQ in *G. oxydans*.

## Conclusions

Bioleaching has the potential to revolutionize the environmental impact of REE production, and dramatically increase access to these critical ingredients for sustainable energy technology. But, making REE bioleaching cost competitive with thermochemical methods will require increasing both the rate and completeness (overall efficiency) of REE extraction. This work gives us a roadmap for improving bioleaching by genetic engineering.

By constructing a whole genome knockout collection for *G. oxydans*, one of the most promising organisms for REE bioleaching, we are able to characterize the genetics of this process with high sensitivity and high completeness. In total we have identified 165 gene disruption mutants that significantly change the acidity of its biolixiviant, rate of production, or both. The regulatory elements of each of these genes represents a dial that can be turned to improve bioleaching.

As well as producing gluconic acid, *G. oxydans* can be used in the production of industrially important products including 2,5-Diketogluconic Acid, a precursor of Vitamin C^48^; 5-Ketogluconic Acid, a precursor of L(+)-tartaric acid used in the production of food and pharmaceuticals^48^; vinegar; sorbitol; and dihydroxyacetone^48^. The *G. oxydans* knockout collection will allow characterization of the production of these metabolites and improvement of their production.

REE bioleaching by *G. oxydans* is predominantly controlled by two well-characterized systems: phosphate signaling and glucose oxidation that is supported by production of the redox cofactor PQQ. Interrupting phosphate signaling control of biolixiviant production by disrupting a single gene (*pstC*) can increase REE extraction by 18%. Disrupting the supply of the PQQ cofactor to the membrane bound glucose dehydrogenase reduces REE extraction by up to 92%.

Comprehensive screening of the *G. oxydans* genome also revealed completely new targets that contribute to REE bioleaching. For example, disrupting *GO_1415*, encoding a protein of unknown function, increases REE bioleaching by 15%. Additionally, our results highlight the potential for a previously uncharacterized role of TldE in PQQ synthesis. Several periplasmic dehydrogenases in *G. oxydans* depend on PQQ for their function^49^, and in the case of the D-sorbitol dehydrogenase, mSLDH, over-expression of the *pqq* synthase genes and *tldD* enhances conversion of N-2-hydroxyethyl-glucamine into 6-(N-hydroxyethyl)-amino-6-deoxy-L-sorbofuranose (6NSL), a precursor to the diabetes drug, Miglitol^50,51^. The discovery of the potential contribution of TldE to PQQ biosynthesis may allow for exceptional enhancement of the cofactor production through the additional over-expression of this gene, and a consequent uptick in dehydrogenase activity. PQQ is an essential cofactor important for several other industrial applications of *G. oxydans*, including production of L-sorbose^52^ and 5-keto-D-gluconate^53^. Furthermore, PQQ alone has many applications across many biological processes from plant protection to neuron regeneration^54^.

Our results are the first ever demonstrating improvement of bioleaching through genetic engineering. Furthermore, the creation of a whole-genome knockout collection in *G. oxydans* will facilitate its use as a model species for further studies in REE bioleaching and other industrially important applications of similar acetic acid bacteria. The findings of the two major systems contributing to acidification in *G. oxydans* suggest the first steps in the roadmap for greatly improving bioleaching: take the brakes off regulation of acid production by disabling the phosphate-specific transport system, while over-expressing *mgdh* along with the expanded synthesis pathway for its cofactor PQQ.

## Materials and Methods

### *Gluconobacter oxydans* B58 Genome Sequencing

Gluconobacter oxydans strain NRRL B-58 (*Go*B58) was obtained from the American Type Culture Collection (ATTC), Manassas, VA. In all experiments, *Go*B58 was cultured in yeast peptone mannitol media (YPM; 5 g L^-1^ yeast extract, 3 g L^-1^ peptone, 25 g L^-1^ mannitol), with or without antibiotic, as specified.

Genomic DNA was extracted from saturated culture using a *Quick*-DNA Miniprep kit from Zymo Research (Part number D3024, Irvine, CA). Genomic DNA library was prepared and sequenced using a TruSeq DNA PCR-Free Library Prep Kit (Illumina, San Diego, CA).

The prepared library was sequenced on a MiSeq Nano (Illumina, San Diego, CA, USA) with a 500 bp kit at the Cornell University Institute of Biotechnology (Ithaca, NY, USA). Resulting paired end reads were trimmed using Trimmomatic^55^ and assembled with SPAdes using k-mer sizes 21, 33, 55, 77, 99, and 127, and auto coverage cutoff^56^. Assembly quality was checked with QUAST^57^ and genome completeness was verified with BUSCO^58^ using the proteobacteria_odb9 database for comparison. The resulting 62 contigs were annotated online using RAST (https://rast.nmpdr.org)^59–61^.

### Gene Ontology Enrichment

DIAMOND^62^ was used to assign annotated protein models with a closest blast hit using the the uniref90 database, an *E*-value threshold of 10^-10^, and a block size of 10. InterProScan^63^ (version 5.50-84.0) was used to assign family and domain information to protein models.

Output from both of these searches was used to assign gene ontologies with BLAST2GO^64^. Gene set enrichment analysis was done with BioConductor topGO package^65^, using the default weight algorithm, the TopGO Fisher test, with a *p*-value threshold of 0.05.

### Mating for Transposon Insertional Mutagenesis

The transposon insertion plasmid, pMiniHimarFRT^39^ was delivered to *Go*B58 by conjugation with *E. coli* WM3064. *E. coli* WM3064 transformed with pMiniHimarFRT was grown overnight to saturation in 50 mL LB (10 g L^-1^ tryptone, 5 g L^-1^ yeast extract, and 10 g L^-1^ NaCl) supplemented with 50 μg mL^-1^ kanamycin (kan) and 90 μM diaminopimelic acid (DAP), rinsed once with 50 mL LB, then re-suspended in 20 mL YPM.

*Go*B58 was grown for approximately 24 hours in YPM, then back-diluted to an optical density (OD) of 0.05 in 750 mL YPM and incubated at 30 °C for two doublings until the OD reached 0.2. *Go*B58 culture was distributed into 13 50 mL conical tubes, to which rinsed and re-suspended WM3064 was added at a ratio of 1:1 by density (approximately 1 mL WM3064 to 50 mL B58). Bacteria were mixed by inversion then spun down at 1900 g for 5 minutes. Supernatant was poured off, and the mixture was resuspended in the remaining liquid (≈ 0.5 mL), pipetted onto a YPM plate in 5 spots of 0.1 mL, and allowed to dry on the bench under a flame.

Mating plates were incubated at 30 °C for 24 hours. Mating spots were collected by adding 4 mL YPM to a plate, scraping the spots into the liquid, then suspending by pipetting up and down several times. Suspended cells were collected from each plate, and the suspension was plated onto YPM agar with 100 μg mL^-1^ kanamycin at 100 μL per plate.

After 3 days of incubation at 30°C, colonies were picked into 96-well microplates using a CP7200 colony picking robot (Norgren Systems, Ronceverte WV, USA). Each well contained 150 μL YPM with 100 μg mL^-1^ kanamycin. For all high-throughput experiments, *Go*B58 was grown in polypropylene microplates sealed with a sterile porous membrane (Aeraseal, Catalog Number BS-25, Excel Scientific) and incubated at 30 °C shaking at 800 rpm. Isolated disruption strains were grown for three days to allow nearly all wells to reach saturation. Wells B2 and E7 of each plate were reserved as no-bacteria controls.

A Monte Carlo numerical simulation (collectionmc^39^) was used to approximate how many insertions would need to occur before a mutant is found representing a knockout of each gene in the genome, which demonstrated that approximately 55,000 mutants would need to be generated and selected to identify mutants in at least 99% of all *Go*B58 genes (**Fig. 1B**).

In total 18 matings were required to recover and pick a progenitor collection of 49,256 disruption strains into 525 microplates over the course of about two months. Microplates with saturated wells were maintained at 4°C for up to 3 weeks and incubated an extra night at 30°C before pooling.

Combinatorial pooling was done in three batches. The 525 plates were virtually arranged in a 20 by 27 grid, and combinatorial pooling, cryopreservation, pool amplicon library generation, and sequencing were all done as previously described^39,40^.

### Curation of a whole-genome knockout collection

Sequencing data for the progenitor collection was processed into a progenitor collection catalog using the KOSUDOKU suite of algorithms^39,40^. To create a condensed collection, a disruption strain was chosen for each of the 2,733 disrupted genes available in the progenitor collection, first prioritizing close proximity to the translation start, then the total probability of the proposed progenitor collection address. A second strain was chosen from the remaining strains for each gene that had another available. For 50 genes, both disruption strains selected were ambiguously located, and thus a third strain was selected from the remaining collection.

In total, 5,137 disruption strains were isolated and struck-out for single colonies. Many progenitor wells were predicted to have more than one possible strain per well, so for each strain, the number of colonies isolated was two times the predicted number of strains in the progenitor well, up to ten. The condensed collection, which amounted to 17,706 wells, was pooled, sequenced, and validated as previously described^39,40^. Unknown disruption strains significantly linked to acidification were identified with Sanger sequencing, also as previously described, with the exception of the transposon-specific primers. For the first and second rounds of nested PCR, the transposon-specific primers were (5’ - GTATCGCCGCTCCCG - 3’, and (5’ - CATCGCCTTCTATCGCCTTC - 3’), respectively.

### Thymol Blue Endpoint Acidity Assay

Endpoint acidity was measured using the pH indicator thymol blue (TB, Sigma-Aldrich, St. Louis, MO), which changes from red to yellow below a pH of 2.8 (https://www.sigmaaldrich.com/US/en/product/sial/114545). The lowest pH of biolixiviant generated by *Go*B58 was 2.3^10^, thus TB allows for distinguishing strains that lower the pH below that of the wild type biolixiviant. To generate biolixiviant, the condensed collection was pin replicated into new growth plates containing 100 μL YPM with 100 μg mL^-1^ kanamycin per well. After two days of growth, an equal volume of 40% w/v glucose was added to the cultures for a final solution of 20% w/v glucose. The amount of glucose needed to lower the pH below 2.3 via the production of gluconic acid was estimated to be 13% w/v, but the higher concentration was used to account for any use of glucose as a carbon source and still maintain an excess amount. Viability tests demonstrated that the bacteria were still viable after two days of culture in such a solution (data not shown).

Bacteria were incubated with glucose for 48 hours to allow acid production to reach completion. Plates were then centrifuged for 3 minutes at 3200 g (top speed) and 90 μL of the biolixiviant supernatant was removed and added to TB at a final concentration of 40 μg mL^-1^. After 1 minute of vortexing, absorbance was measured for each well at 435 nm and 545 nm on a Synergy 2 plate reader (Biotek Instruments, Winooski, VT, USA). Because of variation in background absorbance from well to well on each plate, absorbance was measured at these two wavelengths, and their ratio was used as a proxy for pH, which correlates linearly within the range of pH for the majority of biolixiviants produced by the collection (**Fig. S1**).

### Bromophenol Blue Acidification Rate Screen

Acidification rate was measured using the pH indicating dye, Bromophenol Blue (BPB). Knockout collection strains were grown for two days. OD was measured at 590 nm for each well, then 5 μL of culture was transferred to a polystyrene assay plate containing 95 μL of 2% w/v glucose and 20 μg mL^-1^ BPB in deionized water. The initial pH of the culture is just above 5, and within moments of adding culture to glucose with BPB, the color begins to change rapidly. Assay plates were vortexed for one minute after addition of bacterial culture, then immediately transferred to a plate reader where the change in color was tracked by measuring absorbance at 600 nm every minute for 6 minutes, resulting in 7 reads. Mean rate (*V*) and R-squared were calculated by the Gen5 microplate reader and imager software (Biotek Instruments). A plot of all *V* relative to OD demonstrated that the two are correlated, thus *V* was normalized to OD for each well (**Fig. S3**).

### Hit Identification in Acidification End Point and Rate Screens

Once every well had its assigned data point (A435/A545 for TB, and *V*/OD for BPB), hits were determined by first identifying outliers for each plate. The interquartile range and upper and lower bounds were calculated in Microsoft Excel considering all wells with cultured disruption strains. Any data point that was more than 1.5 times over or under the upper or lower bound, respectively, was considered an outlier. A disruption strain was considered a hit if over half of the wells for that strain (or 1 of 2) were outliers.

### Acidification End Point and Rate Quantification with Colorimetric Dyes

For each assay, knockout strains identified as hits were isolated from the knockout collection into new microplates, along with several blanks per plate, and proxy wild type strains - *Go*B58 strains with an intergenic transposon insertion that should not affect the acidification phenotype (See next methods section). OD and acidification phenotypes were measured for each proxy WT strain separately to verify that growth and acidification are unaffected in these strains (**Fig. S4**).

Acidification phenotypes for the disruption strains (for *n*, see **Tables S6C** and **D**) were compared to that of proxy WT (TB, *n* = 144; BPB, *n* = 31) with a Student’s *t*-test in Microsoft Excel, two-tailed assuming equal variances. A Bonferroni correction was used to determine significance to account for the possibility a comparison is significant by chance alone: a phenotype was considered significant if*p* < 0.05/*N*, where *N* is the number of comparisons being made (*N* = 120 or *N* = 242 for endpoint acidity comparisons with pWT set A or set B, respectively; *N* = 60 for rate of acidification comparisons with pWT).

### Choice of Proxy Wild-type Comparison

The biolixiviant end point pH and acidification rate of each *G. oxydans* mutant were compared against a proxy wild-type set of mutants for each phenotype. To account for the presence of a kanamycin cassette in the genome, the proxy wild-type set for each phenotype was constructed of several mutants with the transposon inserted in an intergenic region, that had no growth defect, and no apparent change in phenotype (**Fig. S2**).

As the efficiency of the *E. coli* WM3064 to *G. oxydans* mating was low, construction of the *G. oxydans* progenitor collection required 18 mating batches. As a result of this, the possibility existed that there might be slight variations in the wild type background from batch to batch.

For the acidificaton rate, these variations did not affect the wild-type behavior across the collection, and a single set of proxy wild-type strains could be used as a comparison with notable disruption strains in the quantification assays. For the end point pH measurement, two distinct proxy wild-type behaviors arose in the condensed collection. Proxy wild-type set A was used for plates 1 to 76; 110 to 130; and 160 to 185, (**Fig. S2A**), and proxy wild-type set B was used for plates 77 to 109 and 130 to 159 (**Fig. S2B**).

Optical density after two days of growth, and endpoint acidity using the TB absorbance ratio (A435/A545) were compared for both wild-type sets (**Fig. S2**). For wild-type set A, which was used for the BPB quantification assay, acidification rate of individual proxy WT strains was also compared. Pairwise comparisons were all made using the emmeans package in R with a Tukey *p*-value adjustment (https://github.com/rvlenth/emmeans).

### Direct Measurement of Biolixiviant pH

Bacteria were grown for 48 hours in tubes containing 4 mL YPM with 100 μg mL^-1^ kanamycin. One tube was left uninoculated as a no-bacteria control. OD was normalized to 1.9 and diluted in half with 40% glucose for a final 20% solution in 1.5 mL. Five replicates were created for each strain and controls, and all mixtures were randomly distributed across two deep well plates. 750 μL of mixture was transferred from each well to a second set of deep-well plates for bioleaching experiments. All plates were incubated shaking at 800 rpm at room temperature.

After two days, one set of deep-well plates was centrifuged for 10 minutes at 3200 g (top speed), and the pH of the supernatant was measured by insertion of a micro-probe to the same depth in each well.

Four standards were used for meter calibration - pH 1, 2, 4, and 7 - and the meter was re-calibrated after every 12 measurements. pH measurements for each disruption strain (n = 5) were compared with those of proxy WT (n = 15) using a Student’s *t*-test in Microsoft Excel, two-tailed, with equal variance. A biolixiviant pH was considered significantly different if*p* < 0.05/N, with N = 22.

### Direct Measurement of REE Bioleaching

The second set of deep-well plates was centrifuged for 10 minutes at 3200 g (top speed), and 500 μL of biolixiviant was transferred from each well to a 1.7 mL Eppendorf tube. 20 mg (4% w/v) of retorted phosphor powder (gift from Idaho National Lab^10^) was added to each tube for bioleaching. Tubes were shaken horizontally for 36 hours at room temperature, then centrifuged to pellet remaining solids. Supernatant with leached REE was filtered through a 0.45 μm AcroPrep Advance 96-well Filter Plates (Pall Corporation, Show Low, AZ, USA) by centrifuging at 1500 × *g* for 5 minutes.

All samples were diluted 1/200 in 2% trace metal grade nitric acid (Thermo Fisher Scientific) and analyzed by an Agilent 7800 ICP-MS for all REE concentrations (*m/z*: Sc, 45; Y, 89; La, 139; Ce, 140; Pr, 141; Nd, 146; Sm, 147; Eu, 153; Gd, 157; Tb, 159; Dy, 163; Ho, 165; Er, 166; Tm, 169; Yb, 172; and Lu, 175) using a rare earth element mix standard (Sigma-Aldrich) and a rhodium in-line internal standard (Sigma-Aldrich, *m/z* = 103). Quality control was performed by periodic measurement of standards, blanks, and repeat samples. A pWT biolixiviant sample without bioleaching was spiked with 100 ppb REE standard and analyzed for all REE concentrations as a control.

An additional 1/20 dilution in 2% nitric acid was analyzed for *δmgdh* and *δpqqc* disruption strains, and the no-bacteria control (glucose).

Bioleaching measurements for each disruption strain (*n* =5) were compared with those of proxy WT (*n* = 15) or glucose (*n* = 5) using a Student’s *t*-test in Microsoft Excel, two-tailed, assuming equal variances. Total REE extracted was considered significantly different if*p* < 0.05/*N*, with *N* = 12 for those compared to pWT, and *N* = 2 for those compared to glucose.

## Data Availability

The datasets generated during and analyzed during the current study are available from the corresponding author (B.B.) on reasonable request.

## Code Availability

The Knockout Sudoku software is available at https://github.com/buzbarstow/kosudoku.

## Materials & Correspondence

Correspondence and material requests should be addressed to B.B.. Individual strains (up to ≈ 10 at a time) are available at no charge for academic researchers. We are happy to supply a duplicate of the entire *G. oxydans* knockout collection to academic researchers, but will require reimbursement for materials, supplies and labor costs. Commercial researchers should contact Cornell Technology Licensing for licensing details.

## Author Contributions

Conceptualization, A.M.S. and B.B.; Methodology, A.M.S. and B.B.; Investigation, A.M.S., B.P., S.M., and B.B; Writing - Original Draft, A.M.S. and B.B.; Writing - Review & Editing, A.M.S., B.P., S.M., M.R., M.W., E.G., and B.B.; Funding Acquisition, A.M.S., E.G., M.W., and B.B.; Resources, M.R., E.G., and B.B.; Supervision, M.R. and B.B.; Data Curation, A.M.S. and B.B.; Visualization, A.M.S. and B.B.; Formal Analysis, A.M.S. and S.M.

## Acknowledgements

We thank D. Reed and Y. Fujita at Idaho National Lab for advice and for gift of rare containing retorted phosphor powder. A.M.S. was supported by a Cornell Energy Systems Institute Postdoctoral Fellowship, and a Small Grant from the Cornell Atkinson Center for Sustainability. This work was supported by Cornell University startup funds, an Academic Venture Fund award from the Atkinson Center for Sustainability at Cornell University, a Career Award at the Scientific Interface from the Burroughs Welcome Fund to B.B., and by ARPA-E award DE-AR0001341 to B.B, E.G., and M.W..

