## Supplementary material for "*Gluconobacter oxydans* Knockout Collection Finds Improved Rare Earth Element Extraction": SI Text

### Supplementary Information Figures

**Fig. S1.** Calibration data for Thymol Blue assay for media acidification.

**Fig. S2.** Comparison of individual proxy WT disruption strains.

### Supplementary Information Tables

**Table S1.** *G. oxydans* B58 genome loci and feature names with transposon insertion sites (see separate Table S1.xlsx)

**Table S2.** Gene Ontology Enrichment Data. (**A**) Non-disrupted genes in *G. oxydans* B58 mutant collection. (**B**) Disrupted genes with lower endpoint acidity. (**C**) Disrupted genes with higher endpoint acidity. (**D**) Disrupted genes with faster acidification. (**E**) Disrupted genes with slower acidification. (See separate Table S2.xlsx).

**Table S3.** Verification of *G. oxydans* B28 mutant identities by Sanger sequencing. (**A**) List of progenitor collection wells verified post-sequencing to confirm the results of location inference. (**B**) List of knockout collection wells verified post-sequencing of the condensed collection to verify strain confirmations. Wells with no "Illumina confirmed transposon coordinate" were unable to be confirmed by Illumina sequencing, but were hits in one of the screens, thus their transposon location was determined by Sanger sequencing.(See separate Table S3.xlsx).

**Table S4.** *G. oxydans* B58 condensed collection catalog. (See separate Table S4.xlsx).

**Table S5.** Results of initial acidification screens of *G. oxydans* B58 knockout collection. (**A**) All hits from Thymol Blue screen for end point pH and Bromophenol Blue (BPB) screen for acidification rate. H indicates that pH of biolixiviant was higher than the proxy wild-type, L indicates that the pH was lower. S indicates that rate of initial acidification was slower than proxy wild-type, F indicates that the rate was was faster. (**B**) Hits from Thymol Blue screen. (**C**) Hits from BPB screen. (**D**) Overlapping hits that appeared in both screens. (See separate Table S5.xlsx).

**Table S6.** Validation results for initial hits identified by acidification screens of *Gluconobacter oxydans* knockout collection. (**A**) All hits. H indicates that pH of biolixiviant was higher than the proxy wild-type, L indicates that the pH was lower. S indicates that rate of initial acidification was slower than proxy wild-type, F indicates that the rate was was faster. (**B**) All significant hits from end point and acidification rate screens. (**C**) Significant hits from TB end point pH screen. (**D**) Significant hits from BPB acidification rate screen. (**E**) Significant hits from TB screen represented in **Fig. 2D**. (**F**) Significant hits from BPB screen represented in **Fig. 2E**. (See separate Table S6.xlsx).

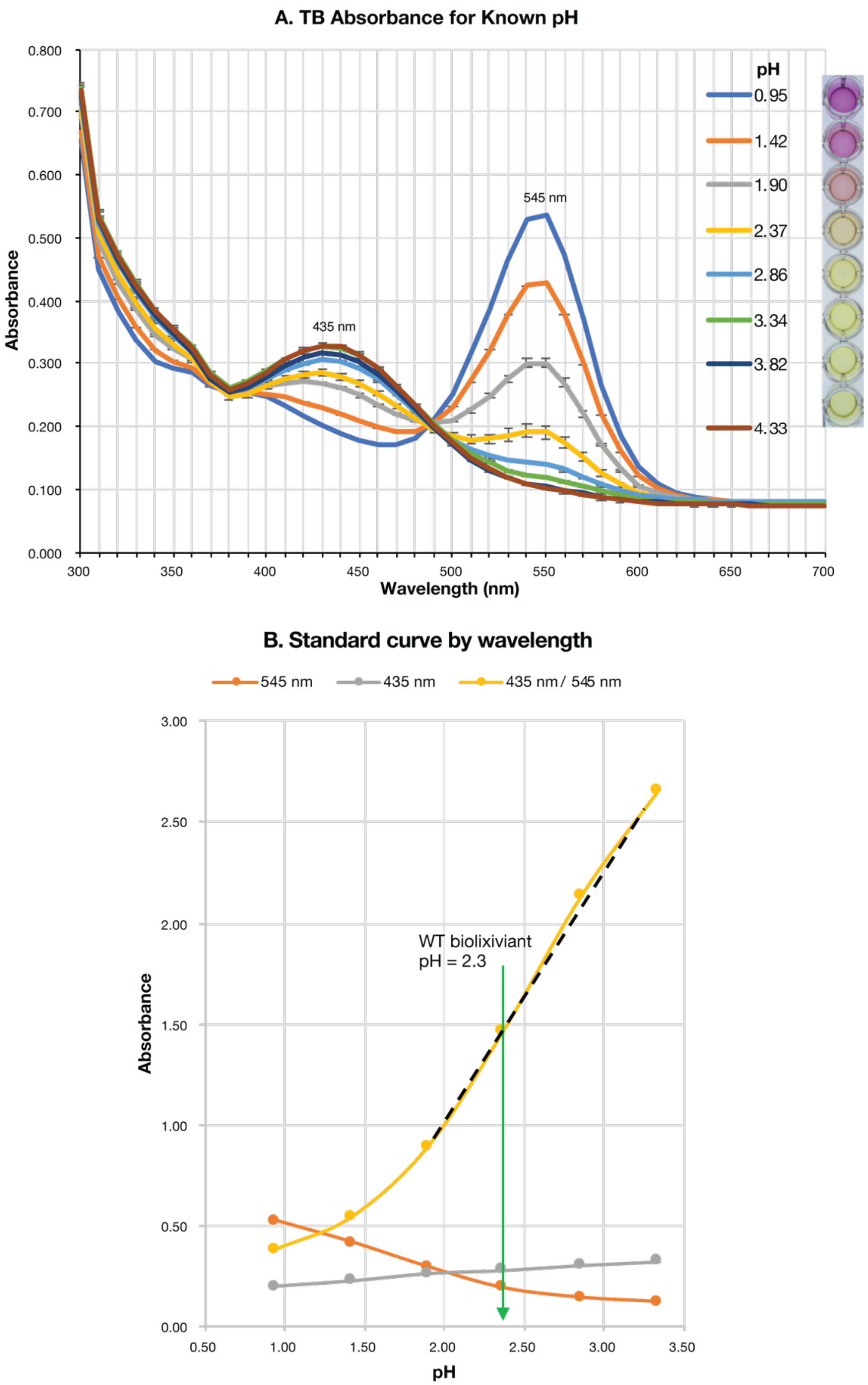


**Figure S1.** Calibration data for Thymol Blue assay for media acidification. (**A**) Thymol Blue (TB) absorbance spectra for solutions of known pH. (**B**) Calibration curve to connect TB absorbance and pH.

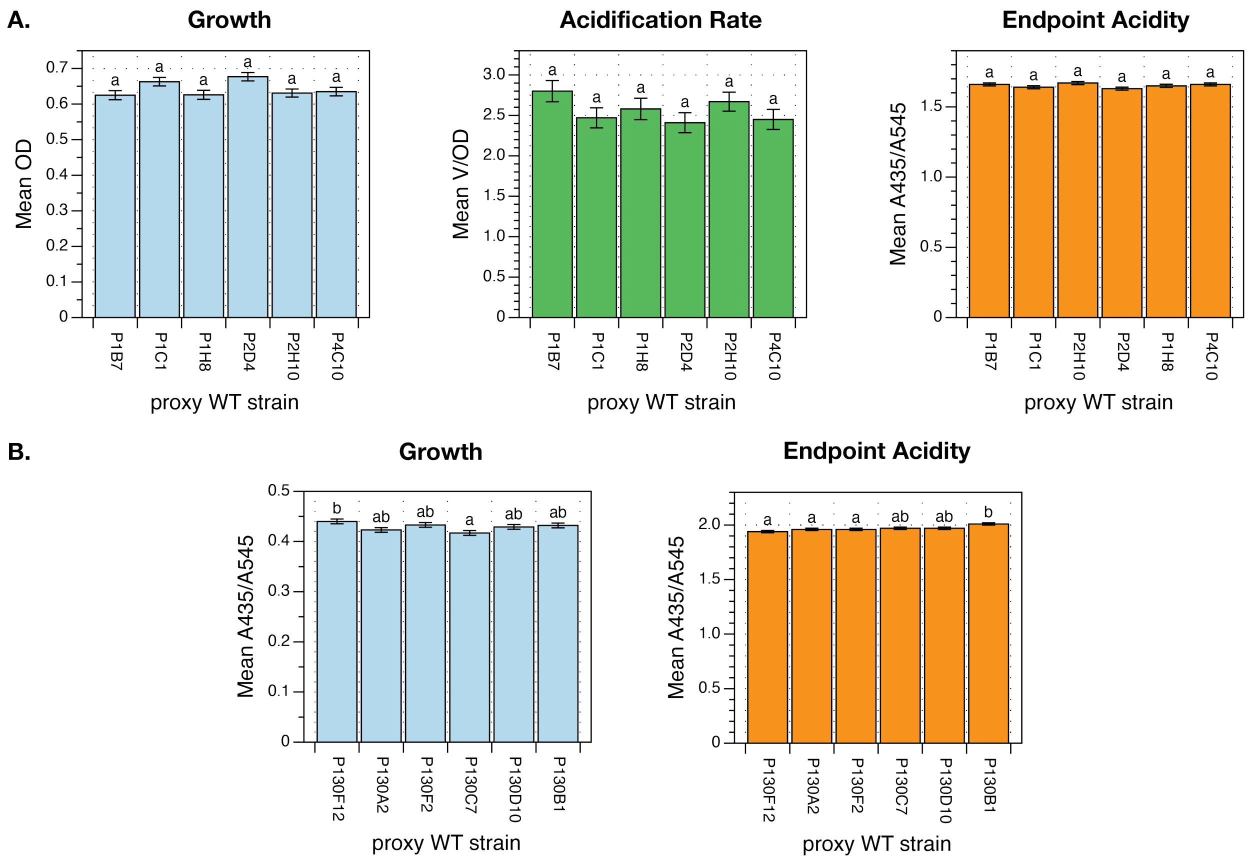


**Figure S2.** Comparison of individual proxy WT disruption strains type A (**A**) and type B (**B**) growth (blue bars), initial acidification rate (green bars), and endpoint acidity (orange bars). Data within each set of proxy WT strains was compared using the emmeans package in R with a Tukey *p*-value adjustment. Letters denote significance groups.
